# Intervertebral Disc Elastography to Relate Shear Modulus and Relaxometry in Compression and Bending

**DOI:** 10.1101/2023.09.01.555817

**Authors:** Zachary R. Davis, Paull C. Gossett, Robert L. Wilson, Woong Kim, Yue Mei, Kent D. Butz, Nancy C. Emery, Eric A. Nauman, Stéphane Avril, Corey P. Neu, Deva D. Chan

## Abstract

Intervertebral disc degeneration is the most recognized cause of low back pain, characterized by the decline of tissue structure and mechanics. Image-based mechanical parameters (e.g., strain, stiffness) may provide an ideal assessment of disc function that is lost with degeneration but unfortunately remains underdeveloped. Moreover, it is unknown whether strain or stiffness of the disc may be predicted by MRI relaxometry (e.g. *T_1_* or *T_2_*), an increasingly accepted quantitative measure of disc structure. In this study, we quantified *T_1_* and *T_2_* relaxation times and in-plane strains using displacement-encoded MRI within the disc under physiological levels of compression and bending. We then estimated shear modulus in orthogonal image planes and compared these values to relaxation times and strains within regions of the disc. Intratissue strain depended on the loading mode, and shear modulus in the nucleus pulposus was typically an order of magnitude lower than the annulus fibrosis, except in bending, where the apparent stiffness depended on the loading. Relative shear moduli estimated from strain data derived under compression generally did not correspond with those from bending experiments, with no correlations in the sagittal plane and only 4 of 15 regions correlated in the coronal plane, suggesting that future inverse models should incorporate multiple loading conditions. Strain imaging and strain-based estimation of material properties may serve as imaging biomarkers to distinguish healthy and diseased discs. Additionally, image-based elastography and relaxometry may be viewed as complementary measures of disc structure and function to assess degeneration in longitudinal studies.

## INTRODUCTION

Low back pain is the leading cause of chronic disability in industrialized Western societies (Murray et al. 2013). Although the causes for low back pain are likely multifactorial, lumbar intervertebral disc degeneration is widely recognized as the most prevalent factor (Endean, Palmer,Coggon 2011). However, the link between degeneration and pain remains controversial because structural indications of degeneration – typically assessed via radiography, computed tomography, and magnetic resonance imaging (MRI) – are also found in asymptomatic individuals (Borenstein et al. 2001). Efforts to resolve this discrepancy are hindered in part by the inherent subjectivity and qualitative nature of these clinical assessments. Furthermore, clinical imaging approaches lack sensitivity to tissue-level changes to composition and mechanical behavior of degenerated disc. Therefore, noninvasive quantification of early disc degeneration remains as a significant challenge.

Disc degeneration, even its earliest form, is characterized by a breakdown of the extracellular matrix (ECM) and a loss of water and proteoglycan content (Antoniou et al. 1996). Because quantitative MRI (qMRI) can relate changes in relaxation time (e.g. *T_1_*, *T_2_*) with alterations to the water, proteoglycan, and collagen content (Chatani et al. 1993; Weidenbaum et al. 1992), relaxometry biomarkers can be used to evaluate the extent of disc degeneration. *T_1_* and *T_2_* have been correlated to degeneration grade(Antoniou et al. 1998; Marinelli, Haughton, Anderson 2010) and tissue macromolecule composition, including proteoglycan (Marinelli et al. 2009) and collagen (Antoniou et al. 2006). Furthermore, relaxation times indicate a spatial distribution of biochemical content, enabling localization of tissue degeneration (Ellingson et al. 2013). However, because relaxation time also depends on factors such as collagen orientation (Xia et al. 1997), age (Marinelli, Haughton,Anderson 2010), and mechanical loading history (Chiu et al. 2001), the interpretation of relaxometry is complex and may not directly correlate to the mechanical behavior of a disc under load. Assessments of mechanical function like intratissue strains may provide independent and complimentary imaging biomarkers for evaluation of disc degeneration.

MRI has additionally been used for full-field strain measurement of the disc under mechanical loading. MRI-based approaches to calculate tissue deformation and strain in the disc include warp field image registration (Yoder et al. 2014), digital image correlation (O’Connell et al. 2007; O’Connell et al. 2011; O’Connell, Vresilovic, Elliott 2011; Tavana et al. 2020), and displacements measured under applied loading by MRI (dualMRI) (Chan and Neu 2014; Wilson et al. 2021). dualMRI has been used previously to characterize strain behavior in cartilage and intervertebral disc (Chan et al. 2011; Chan, Neu,Hull 2009; Griebel et al. 2014; Neu and Walton 2008) and can be readily adapted for measurement of *in vivo* tissue deformations (Chan et al. 2016; Wilson et al. 2021).

Full field displacements and strains, such as those derived from dualMRI, also enable the estimation of elastography, or spatial maps of displacements, strain, or material properties (e.g. shear modulus). *Ex vivo* and *in vivo* MRI elastography (MRE) has produced multi-dimensional shear modulus maps of discs (Cortes et al. 2014; Muthupillai et al. 1995; Streitberger et al. 2015; Walter et al. 2017), this method is generally limited to high frequency (∼1000 Hz) shear waves that may not reflect the properties of the viscoelastic disc under normal (low frequency; ∼1 Hz) activities like walking or bending. In contrast, dualMRI synchronizes mechanical loading at more physiologically relevant loading frequencies (i.e., 1-2 orders of magnitude lower than MRE) to cyclic phase-contrast image acquisition.

Whereas tissue content and material properties dictate but are not altered by loading, the mechanical behavior of the disc is inextricably linked to the applied loading and other boundary conditions. Inverse modelling can be used to estimate mechanical parameters using image-based, full-field strain (Avril et al. 2008a). These inverse modeling approaches have been successful in estimating stress tensors (Kim et al. 2012), stiffness ratio (Avril et al. 2008b), and shear moduli (Avril et al. 2008b). Recent developments have improved the accuracy of calculating shear modulus of soft tissue-like materials (Mei and Avril 2019). Therefore, our objectives in this study were twofold: (1) to implement inverse modeling for the estimation of shear moduli from dualMRI of human cadaveric disc, and (2) to investigate the correlation of mechanical parameters to MRI-based biomarkers associated with tissue composition (*T_1_* and *T_2_*) as functions of region within the disc.

## MATERIALS AND METHODS

### Specimen Preparation

Human lumbar L4-L5 motion segments from three donors (1 female and 2 male, 35±13yrs, range: 22-48yrs, height: 172±12cm, weight: 92±17 kg) were procured (Unyts, Buffalo, NY). The vertebral bodies were isolated by transecting the vertebrae at the pedicles. Excess tissues, including the pedicles, laminae, superior and inferior articular processes, and the transverse and spinous processes, were removed while preserving the anterior and posterior longitudinal ligaments and the intervertebral disc. The samples were visually inspected during dissection to check for features indicative of disc degeneration (i.e., concise margins between the vertebral bodies and the discs, ample disc space, lack of bony spurs).

L4 and L5 vertebral bodies were secured using fiberglass resin into a sample holder, which was connected to an electro-pneumatic loading system compatible with a 9.4-Tesla horizontal bore MRI system (Bruker GMBH, Ettlingen, Germany; Figure 1A) (Chan et al. 2014).The design of the sample rig allowed for a quick interchange of loading modes from axial compression to bending mode by the removal of a support pin (Figure 1B). To prevent desiccation, the tissues were wrapped with PBS-soaked gauze throughout the cyclic loading and MR imaging experiment, and PBS was replenished as needed.

**Figure 1.**
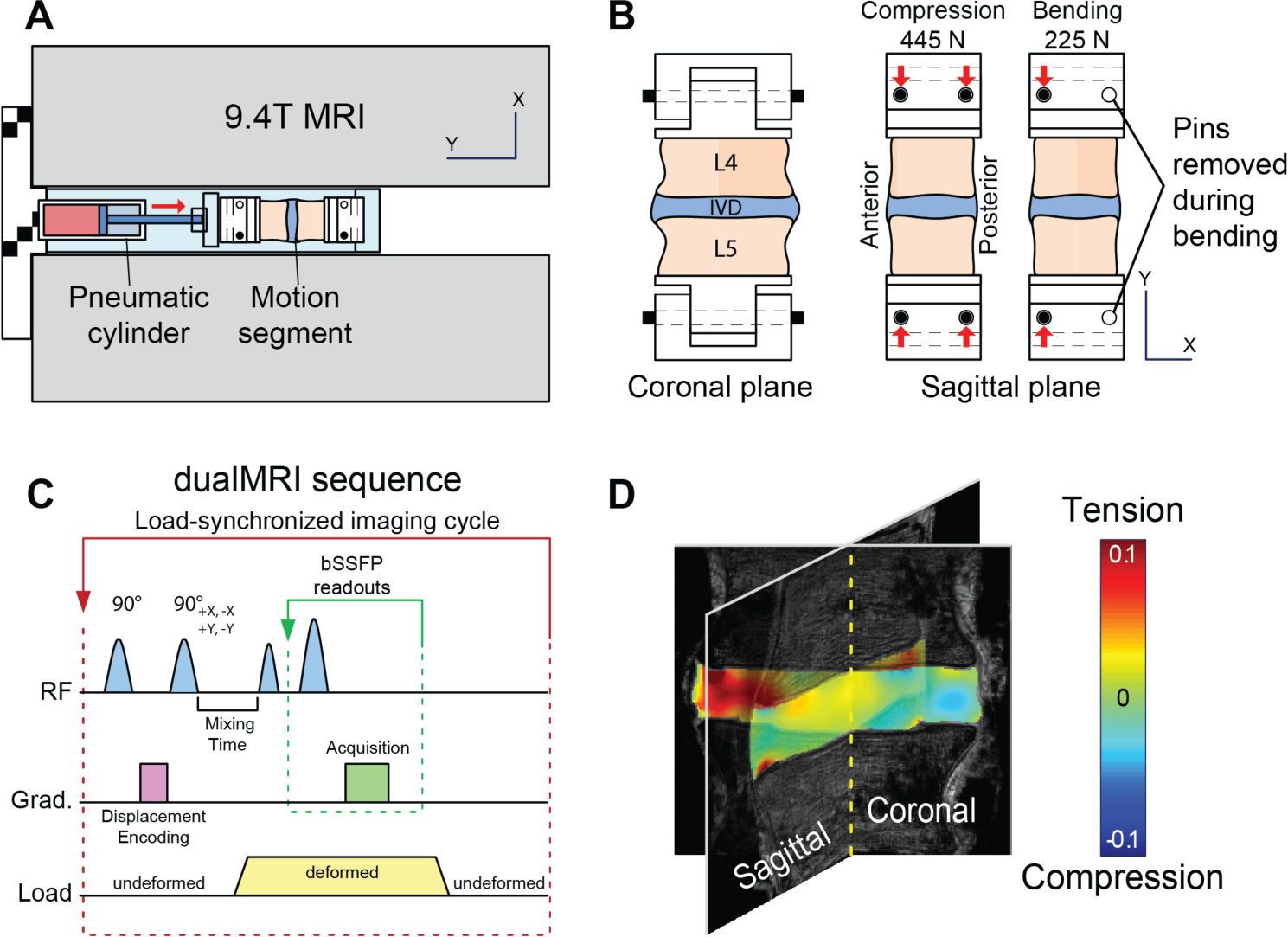
Experimental setup for MRI-based elastography measurements throughout the interior of the intervertebral disc (IVD) in compression and bending. (a) MRI-compatible electro-pneumatic loading system designed for a 9.4-Tesla MRI system. (b) Support pin configuration permits toggle of loading modes from centric to eccentric axial compression, the latter of which generates bending loads. (c) Loading profile synchronized with displacement encoding with stimulated echoes (DENSE) for dualMRI.

### T_1_ and T_2_ Mapping

Prior to loading, *T_1_* and *T_2_* mapping of the disc was performed in both sagittal and coronal planes, approximately through centroid of the disc (Figure 2). Scan parameters were field of view = 64×64mm^2^, spatial resolution = 250×250µm^2^, slice thickness = 2mm. For *T_1_* relaxation time mapping, a fast spin echo acquisition was used with multiple repetition times (TR = 100, 300, 500, 1000, 2000, 4000 ms) and an echo time (TE) of 10ms. For *T_2_* relaxation time mapping, fast spin echo acquisition parameters were TE = 20, 60, 100, 141, 181, 221, 261, 301 ms and TR = 4000 ms. Image analysis software (Paravision, Bruker GMBH, Ettlington, Germany) was used to estimate *T_1_* and *T_2_* at each pixel of interest with exponential fitting.

**Figure 2.**
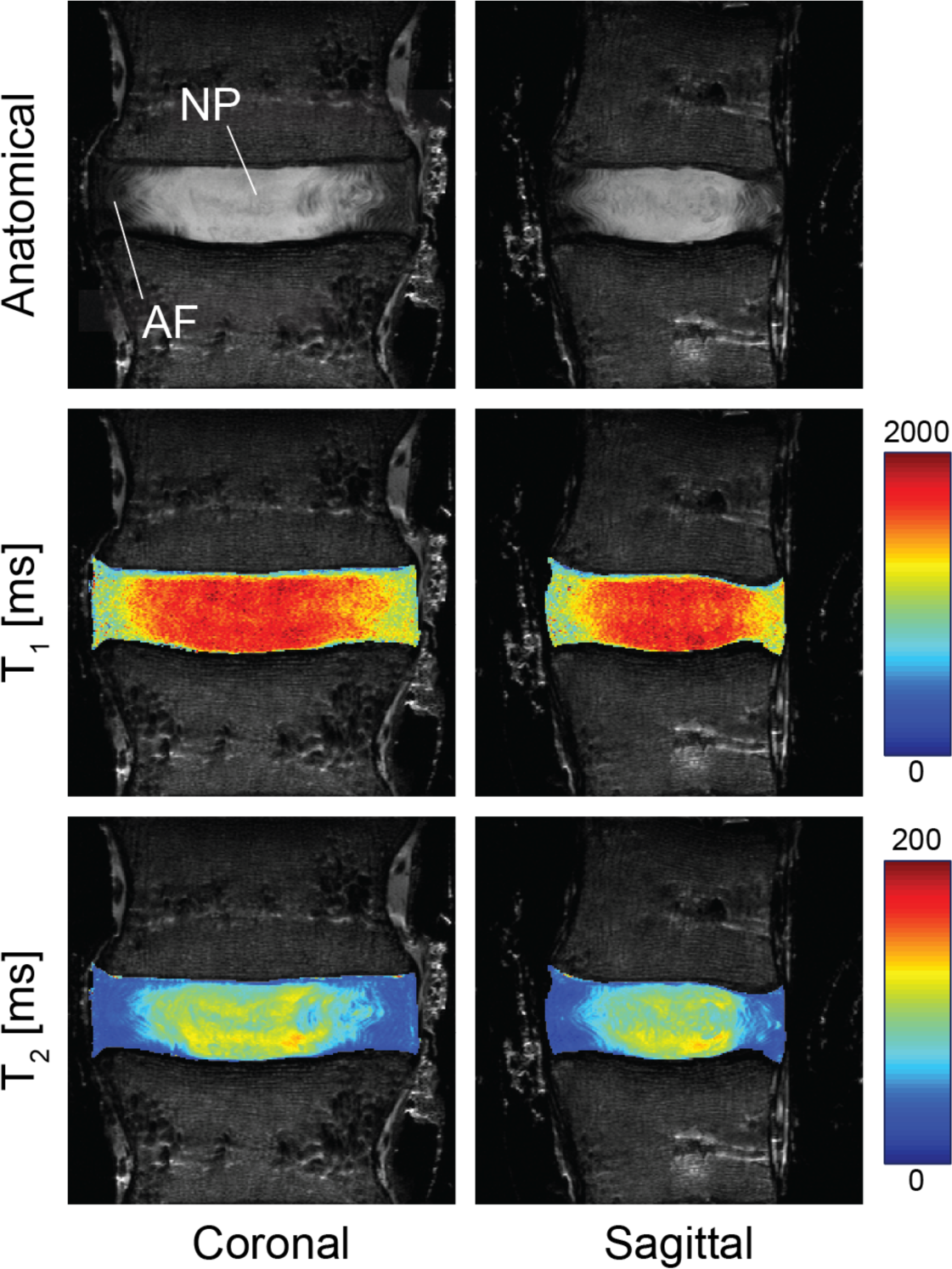
Representative morphometric and T1/T2 relaxation time maps in coronal and sagittal imaging planes. Morphometric (proton density-weighted) images indicate a bright, fluid-rich nucleus pulposus surrounded by the annulus fibrosis. Elevated values for *T_1_* and *T_2_* are observed in the nucleus pulposus compared to the annulus fibrosis, corresponding the fluid levels in individual tissue compartments. Relaxation time maps suggest potential imaging-based biomarkers that could also serve as surrogates for biomechanical function.

### dualMRI and Strain Mapping

Using dualMRI, 2D Green-Lagrange strains (*E_xx_*, *E_yy_*, *E_xy_*) were measured within coronal and sagittal imaging planes under cyclic compression and bending. In the axial compression mode, 445 N was applied along the superoinferior axis to simulate force experienced during normal gait (Cappozzo 1984; Rohlmann et al. 2014). In the bending mode, the posterior support pins were removed to create a 1.33-cm offset to the applied compression (225 N, Figure 1B). This created a 3.0-N·m bending moment in the anterior direction, which is a typical magnitude within the lumbar spine under non-strenuous movements (Adams and Dolan 1991; Rohlmann et al. 2014). Loading was held for 2 seconds to permit MRI imaging and followed by a 3-second unloaded recovery (Figure 1C). Preconditioning cycles were applied for 30 min to achieve a steady-state response and minimize motion artifacts before the start of load-synchronized imaging (Chan and Neu 2012).

For dualMRI (Figure 1C), displacements were encoded at 0.32 rad/mm (Chan and Neu 2012), and phase cycling was used to eliminate artefacts (Epstein and Gilson 2004). Acquisition of images, using the same resolution and field of view as *T_1_* and *T_2_* mapping, was accomplished with balanced steady state free precession (bSSFP, TE/TR=1.607ms/3.215ms, flip angle=25°). Custom scripts (Matlab R2012a, The MathWorks Inc., Natick, MA) was used to calculate displacements and strains (Chan and Neu 2012). Strains were calculated with respect to the imaging coordinate system (*E_xx_*, *E_yy_*, *E_xy_*), and principal strains (*E_1_, E_2_*) and maximum shear strain (*γ_max_*) were also determined.

### Inverse Modeling

We estimated the shear modulus of each disc utilizing an iterative inverse approach, minimizing the gap between measured and computed displacement fields throughout the region of interest in L2 norm (Oberai, Gokhale,Feij o 2003). The computed displacement field was obtained by solving the forward problem using the finite element method. In the forward model, we assumed that the disc satisfies an incompressible linear elastic constitutive behavior. In the incompressible and linear elastic model, only the shear modulus needs to be determined. Since tissue displacement was measured only within single image planes through the disc, we also assumed that the disc was in a 2D plane strain state. In the finite element forward simulation, we use stabilized finite elements to address volumetric locking related to incompressibility. In the inverse modeling, we introduced a regularization term in the cost function to smooth the reconstructed elastic property distribution and avoid overfitting. The optimization problem was solved by the quasi-Newton method where the derivative of the objective function with respect to every shear modulus in the domain of interest is required. To reduce the computation cost, an adjoint-based method was employed. This iterative process terminated when the difference of the objective function values or the associated gradients between two neighboring iterations were less a threshold. These procedures were used to estimate relative shear modulus from the image-based strains that resulted from cyclic compression and bending.

### Image and Statistical Analysis

Displacement maps from dualMRI were used to map *T_1_* and *T_2_*, which were measured in the undeformed disc, onto the deformed disc geometry, the image space within which strains were calculated and moduli estimated. Relaxation times (*T_1_, T_2_*) and mechanical parameters (*G*, 2D strains) were evaluated for all discs using MATLAB. Data are reported as mean ± standard deviation across the entire region of interest. Paired t tests were used to compare average shear moduli estimated from compression or bending. Distributions of these parameters were also visualized for each disc using histograms for qualitative comparison.

To evaluate the relationships among different relaxation time and mechanical parameters, regional analyses were performed. Correlations between relative shear modulus as estimated from compression (*G_(c)_*) and relative shear modulus as estimated from bending experiments (*G_(b)_*) were calculated. Correlations between relaxation-time maps (*T_1_, T_2_*) and relative shear moduli (*G_(c)_*, *G_(b)_*) were separately evaluated. Each disc was divided into five evenly spaced regions horizontally and three regions vertically (top 25%, middle 50%, and bottom 25% by height), resulting in a total of 15 regions in each disc. These divisions followed the contours of the disc region of interest. Within each of these regions, Pearson’s correlation was used to evaluate the relationship between each pair of parameters. Statistical significance was defined at α = 0.05 for all tests.

## RESULTS

### T_1_ and T_2_ Relaxation Time Mapping

*T_1_* relaxation time from coronal and sagittal planes were 1300 ± 265 and 1308 ± 270 ms, and *T_2_* were 71 ± 29 and 71 ± 28 ms, respectively, taken across the full disc. Regions of elevated *T_1_* and *T_2_* values were observed in the central region of the disc in both coronal and sagittal planes (Figure 2), suggesting the location and margins of the NP and AF.

### dualMRI and Strain Mapping

Strain fields, measured under compression and bending using dualMRI, were heterogeneous in both coronal and sagittal planes (Figures 3 and 4). Mean strains *E_xx_*, *E_yy_* and *E_xy_* were 0.018 ± 0.006, −0.031 ± 0.006 and 0.002 ± 0.002, respectively, taken across all discs (Table 1). Average principal strain measures (*E_1_*, *E_2_*, *γ_max_*) were 0.0166 ± 0.0075, −0.0510 ± 0.0045, and 0.0338 ± 0.0023, respectively, across all discs. Under compression, the location of maximum *E_xx_* and *E_yy_* within the disc showed no apparent pattern in neither coronal nor sagittal planes among the discs; however, under bending, locations of strain maxima appeared to be more consistent. In the coronal plane, maxima were located at the midline of the disc, and, in the sagittal plane, at the posterior aspect. A significant difference between compression and bending was found in first principal strain (p = 0.029) and maximum shear stress (0.013) calculated in the coronal plane but for no other mechanical parameters nor in the sagittal plane.

**Figure 3.**
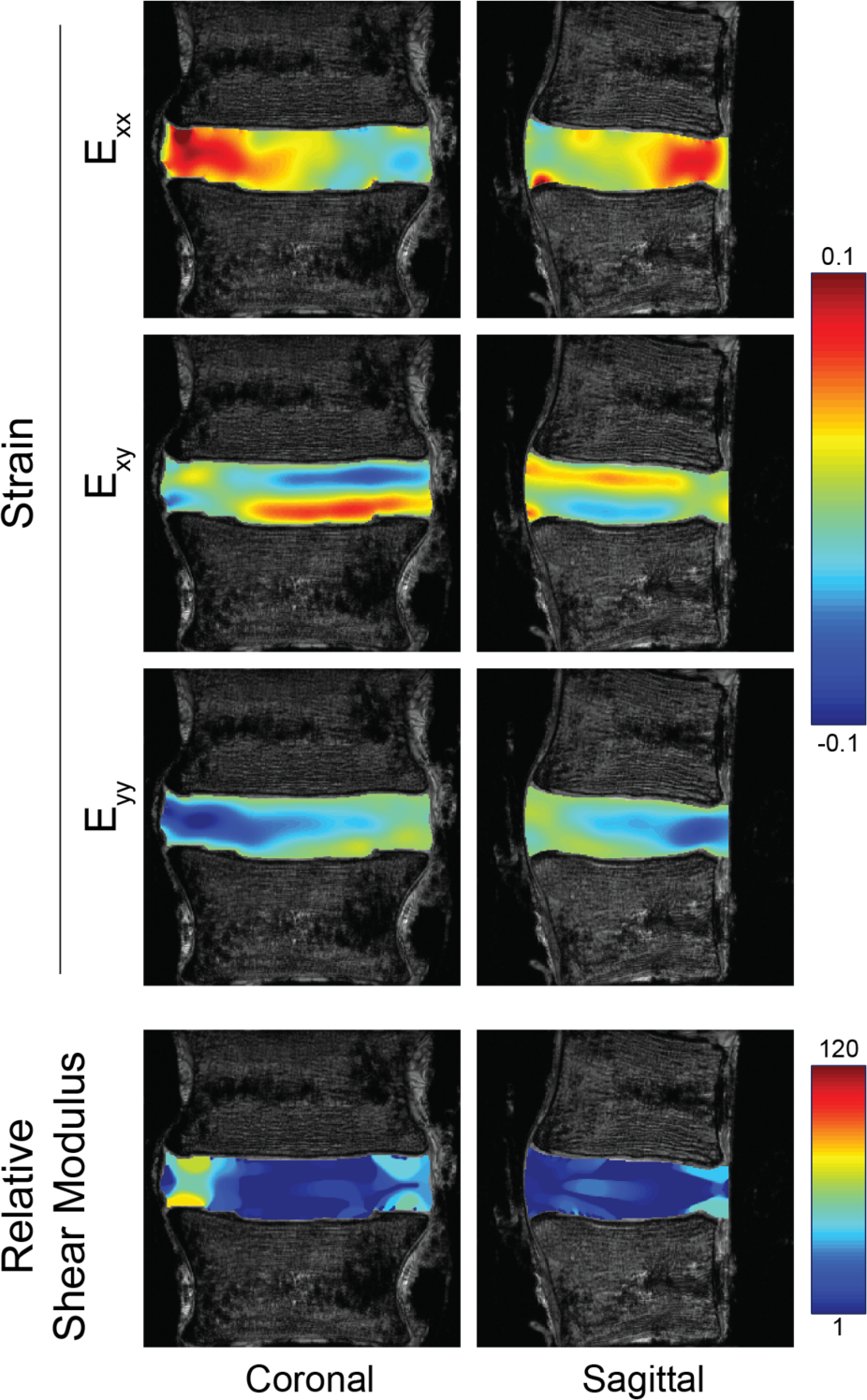
Representative Green-Lagrange Strains and Corresponding Shear Modulus Maps under Compression. Strains in coronal and sagittal image planes were calculated from displacement maps derived from displacement-encoded MRI under cyclic axial compression. Relative shear modulus was calculated separately using strain data from each image plane. Strains and moduli are shown for a representative specimen.

**Figure 4.**
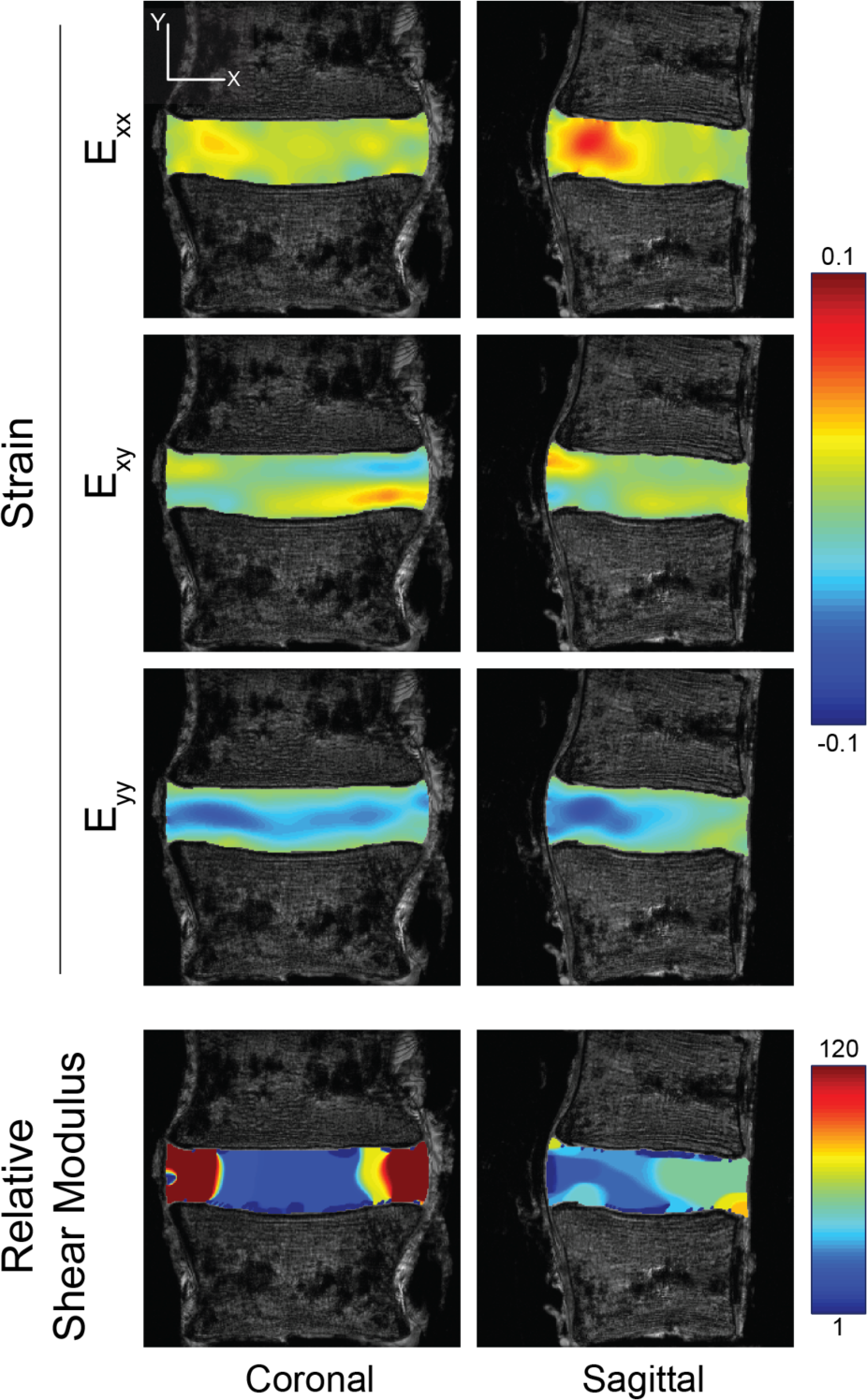
Representative Green-Lagrange Strains and Corresponding Shear Modulus Maps under Bending. Strains in coronal and sagittal image planes were calculated from displacement maps derived from displacement-encoded MRI under cyclic bending. Relative shear modulus was calculated separately using strain data from each image plane. Strains and moduli are shown for a representative specimen.

**Table 1.**
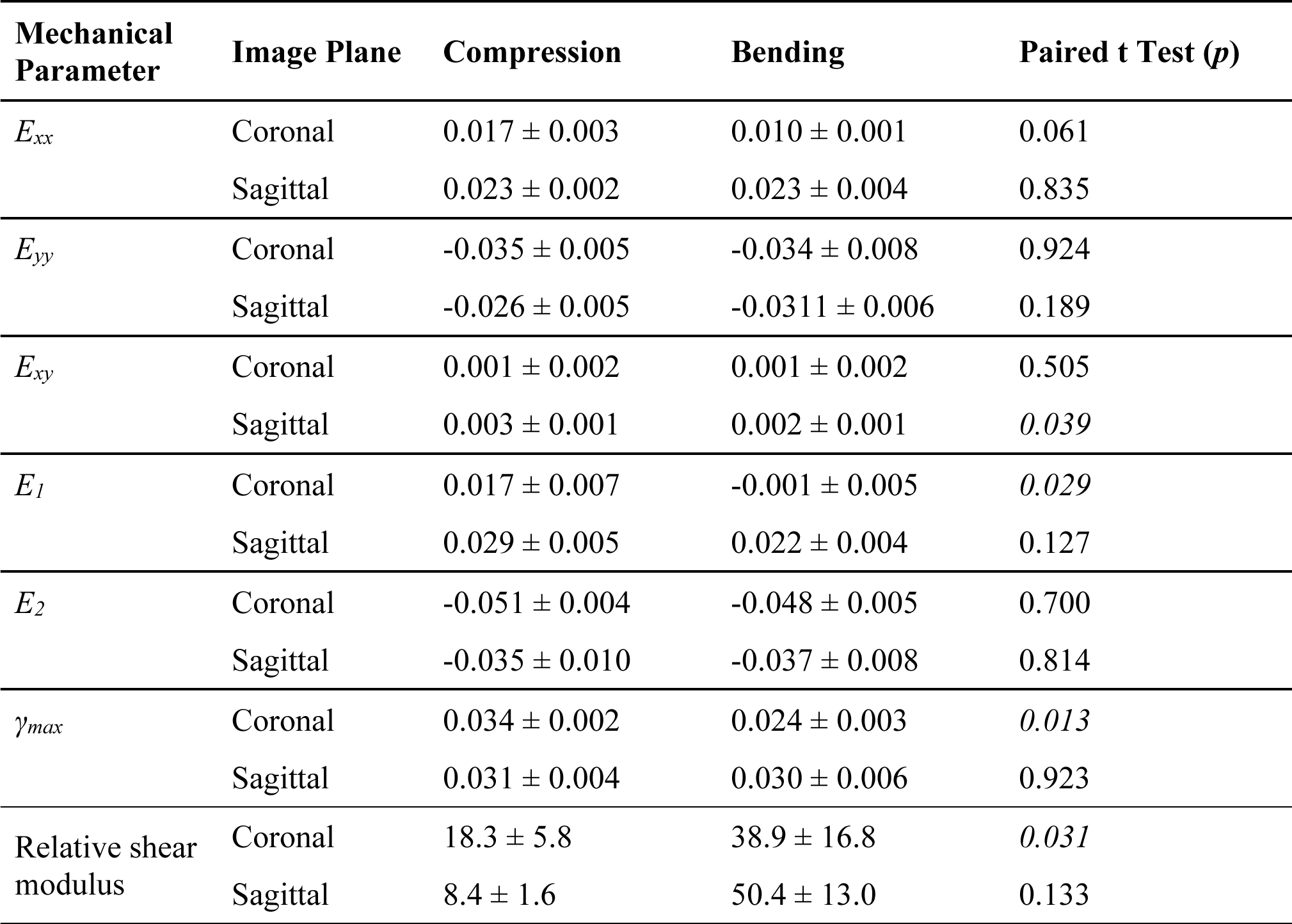
Average Mechanical Parameters across all Discs. In-plane strain calculated from displacement-encoded imaging (dualMRI) and relative shear moduli were averaged over the entire disc regions of interest in the indicated image planes. Strains were calculated with respect to the image plane (*E_xx_*, *E_yy_*, *E_xy_*) as well as in in-plane principal directions (*E_1_*, *E_2_*). Maximum shear strain (*γ_max_*) was also determined per pixel. Data represents mean ± standard deviation (*n*=3 biological replicates). Significant differences in a mechanical parameter estimated from compression vs. estimated from bending experiments are indicated in bolded *p* values.

### Shear Modulus

Estimates of shear modulus from compression-only testing (Table 1) demonstrated relatively stiff AFs and compliant NPs (Figure 3, lower panels). However, discs showed an apparent stiffening in both the AFs and anterior aspects of the NP under bending (Figure 4, lower panels). Relative shear moduli estimated from strain maps taken under bending (Table 1) were significantly greater than moduli estimated with compression in the sagittal (p = 0.031) but not coronal (p = 0.133) planes.

### Comparison of Relaxometry and Mechanical Parameters

The relationships between shear modulus and relaxation times were evaluated by region in the coronal (Figure 5) and sagittal images (Figure 6). Each disc in the coronal and sagittal planes was divided into 15 regions for analysis (Figures 5A and 6A). The distribution of relaxation times and relative shear moduli was evaluated across all discs (Figures 5B-C and 6B-C). Interestingly, relative shear modulus estimates derived from compression and bending experiments were better correlated – primarily in the regions that correspond to the anterior inner AF and outer NP – in the coronal plane than the sagittal plane (Figures 5D and 6D).

**Figure 5.**
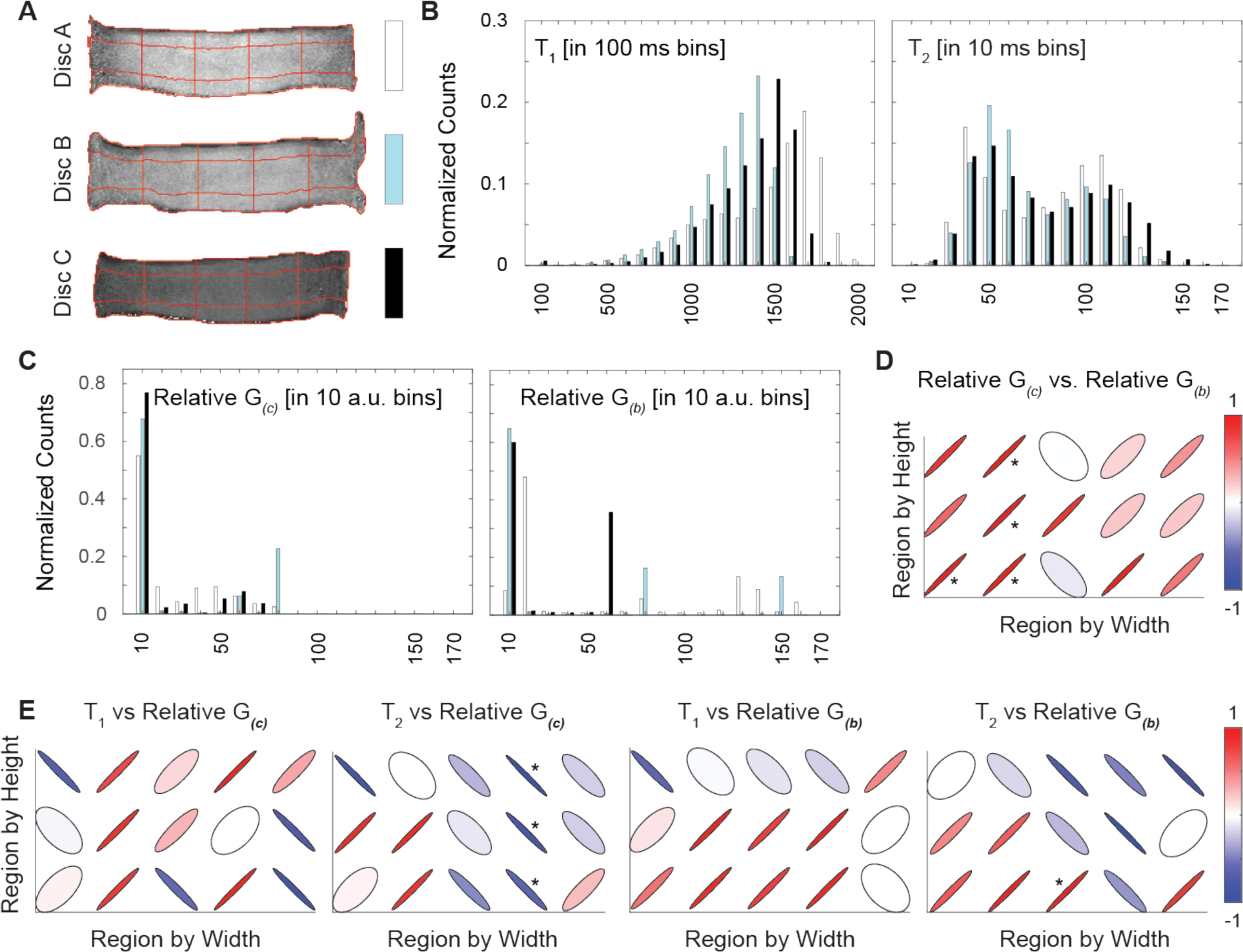
Relaxation Time and Relative Shear Moduli in the Coronal Plane. Each disc was segmented into 15 regions (A) for analysis of parameters in the coronal image plane. Histograms (with each disc indicated with white, aqua, or black bars) of the relaxation times (*T_1_*, *T_2_*) and relative shear moduli estimated in compression and bending (*G_(c)_*, *G_(b)_*) showed no qualitative differences among discs (B, C). Correlations between *G_(c)_* and *G_(b)_* and between relaxation times and shear moduli were calculated within each region (E) (* p < 0.05). For correlations, the color bars represent the sign and R^2^ value of the correlations in each region.

**Figure 6.**
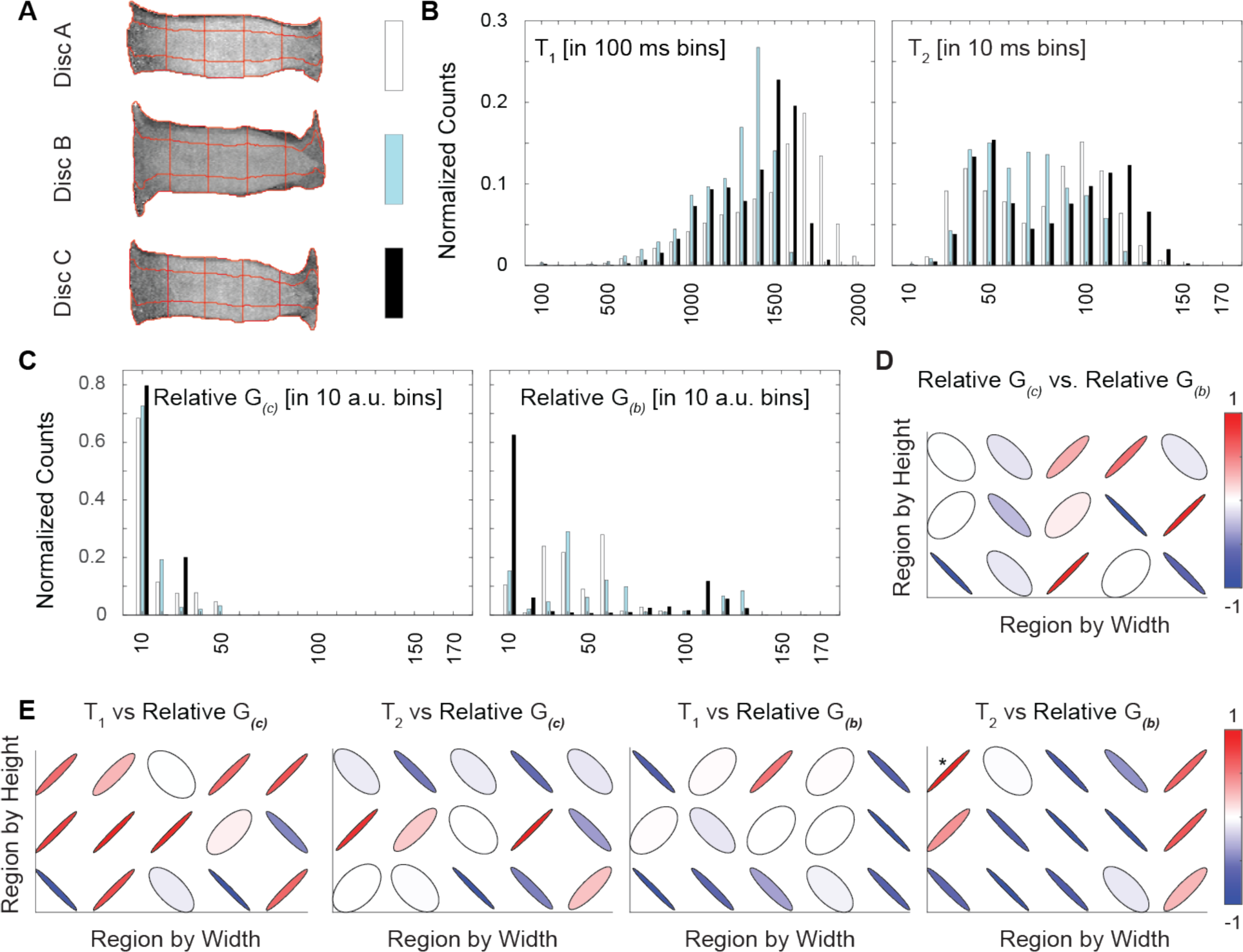
Relaxation Time and Relative Shear Moduli in the Sagittal Plane. Each disc was segmented into 15 regions (A) for analysis of parameters in the sagittal image plane. Histograms (with each disc indicated with white, aqua, or black bars) of the relaxation times (*T_1_*, *T_2_*) and relative shear moduli estimated in compression and bending (*G_(c)_*, *G_(b)_*) showed no qualitative differences among discs (B, C). Correlations between *G_(c)_* and *G_(b)_* and between relaxation times and shear moduli were calculated within each region (E) (* p < 0.05). For correlations, the color bars represent the sign and R^2^ value of the correlations in each region.

Under compression, relative shear modulus showed significant correlations with relaxation times in four regions in the coronal plane but none in the sagittal plane (Figures 5E and 6E). Under bending, relative shear modulus showed significant correlations with relaxation times in only one region (Figures 5E and 6E). Of the 15 total regions analyzed, ten did not show any significant correlations in any combination of relaxation time and shear modulus. Of the significant correlations, only one was between *T_1_* and relative shear modulus, while *T_2_* and relative shear modulus were correlated in six regions. Under compression in both the sagittal plane and coronal plane, few statistically significant correlations were found between strains and relaxation times in any of the 15 regions within the discs (Supplemental Figures 1 & 2). Under bending, four regions in the sagittal plane and only one in the coronal plane had statistically significant correlations between strains and relaxation times.

## DISCUSSION

In this study we used dualMRI in orthogonal (coronal and sagittal) imaging planes to evaluate in-plane strains resulting from cyclic compression and axial bending in human lumbar intervertebral discs. Futhermore, to investigate the utility of MRI relaxometry in the disc as a surrogate parameter to assess mechanical function of the tissue under load, we examined correlations among relaxation times (*T_1_*, *T_2_*), relative shear modulus, and principal strains (*E_1_*, *E_2_*, *γ_max_*). Average strain measures do not capture the complexity and heterogeneity of strains observed throughout the tissue, which motivates the need for spatial mapping of mechanical behavior. Moreover, full-field displacement and strain data enable quantification of spatial maps of material properties and elastography as new potential biomarkers for IVD health. Although fully three dimensional dualMRI would suffer from impracticably long imaging times, doubling the imaging time by acquiring tissue strains in orthogonal directions could be a promising approach to obtaining greater information for inverse modeling. The goals of this study were (1) to evaluate MRI relaxation times and dualMRI-derived mechanical parameters such as strain and estimated modulus in orthogonal anatomic planes under two common modes of loading in the disc and (2) to determine if MRI relaxation times and mechanical parameters can act as mutual surrogates in characterizing heterogeneity within the disc.

The mechanical function of the disc is closely associated with the structure and content of the extracellular matrix (ECM) within both the nucleus pulposus (NP) and annulus fibrosis (AF) (Inoue and Espinoza Orias 2011). Therefore, direct measurement of the mechanical behavior of a disc under load could provide a more comprehensive picture of disc health, including the physical origin of pain, than structure or content biomarkers alone. In prior studies, measurement of nominal changes in disc height via MRI (Dimitriadis et al. 2012), video fluoroscopy (Nagel et al. 2014), ultrasound (Zheng et al. 2014), and dynamic radiographic imaging (Byrne, Aiyangar,Zhang 2019) under extension and flexion have been used to estimate disc deformation *in vivo*. However, these methods are often based on nominal measures (e.g. distances between endplate surfaces) and do not permit measurement of internal mechanics required to detect the focal and heterogeneous changes with disc degeneration (Boos et al. 2002).

Interestingly, deformable image registration methods (e.g., warp field, digital image correlation) have been used to calcluate strain in the AF using tracking of intrinsic textural features visible in the morphological images (O’Connell et al. 2007; Yoder et al. 2014). In contrast to texture correlation, the phase encoded data in dualMRI enables the measurement of tissue displacements with high precision and resolution independent of image texture (Chan and Neu 2014), an advantage for disc imaging because the NP may lack readily trackable features or textures.

While dualMRI experiments can be designed to understand cyclic processes that mimic normal activities like walking, a caveat of this technique is that it cannot measure the immediate mechanical response of tissue in its fully hydrated and undeformed state (e.g., strains under a single impact load). Despite this limitation, dualMRI remains a highly precise technique for the direct measurement of deformations in tissues, which, under normal conditions, are more often intermittently loaded than not. The same regions in different samples sometimes resulted in a variance for shear modulus. For example, the same central region under bending in the sagittal plane had an average relative shear modulus of 27.45 ± 23.15 across the three samples. There are multiple factors that could cause the high variation, including differences among donors and any image noise that would increase the expected error from the inverse method. Furthermore, the inverse method assumed that each imaged cross section of the IVD was in a state of plane strain. Neglecting strains in the out-of-plane direction could induce artefacts in the obtained shear moduli, with variations from an individual to another depending on the subject specific morphology of each IVD. Another limitation to this study is that there were only three samples used for the correlations with little background on previous health history. Finding multiple samples of a similar age and health history would allow for a better understanding of MRI relaxometry and mechanical property correlations.

Although relaxometry may provide a straightforward measurement of disc composition, it is unknown whether relaxation time can act as a sole indicator of the mechanical behavior of the disc. Such an indirect link may be reasonable to assume because the qMRI reflects biochemical content and structure of the disc and would naturally influence the mechanical response of the tissue under load. qMRI of the disc may thus complement or support deformation patterns observed under load, as has been previously demonstrated with qMRI and dualMRI in articular cartilage (Griebel et al. 2014). However, more recent work compared principal strains to T_1ρ_ and T_2_ relaxation times within cartilage *in vivo* but found no statistically significant correlations (Wilson et al. 2021).

Based on the regional correlation analysis, we found limited evidence supporting a relationship between mechanical parameters and relaxation times *T_1_* and *T_2_*. Among the correlations with relative shear modulus, only one region was statistically significant between *T_1_* and *G_(c)_* (coronal plane, p = 0.043). Previous studies have shown that relaxometry may be associated with the amount of deformation under compressive loading (O’Connell, Vresilovic, Elliott 2011) and bending stiffness (Ellingson et al. 2013) in human disc and that these values do not drift significantly during or after application of moderate cyclic loading (Chan and Neu 2014). The lack of correlations between *T_2_* and shear modulus is not consistent with previous work within the NP (Cortes et al. 2014), wherein MRE was applied to 16 cadaveric samples. However, in that prior work, inverse mechanics was applied solely to the NP. Other studies have shown inconsistent correlations between *T_1ρ_* relaxation time and radial and axial strains in the AF (O’Connell, Vresilovic, Elliott 2011). The lack of significant correlations in multiple regions of the disc shows that, currently, MRI relaxometry outputs cannot be used as direct proxies for localized mechanical parameters of a tissue, nor vice versa. These results may also suggest that both relaxation times, and potential any other MRI-based correlate to tissue composition or ultrastructure, and mechanics should be assessed independently, consistent with a previous study that correlated strain to relaxation times in the cervical spine (Wilson et al. 2021).

Finally, in comparing the relative shear moduli estimated with strains under compression and bending, we found correlations in the coronal plane but not the sagittal. Since sagittal in-plane strains differ between the two loading regimes, this lack of correlation in estimated shear modulus is not unexpected. The load boundary conditions used in the inverse model provide an inherently differing set of inputs. In the coronal plane, where in-plane strains were expected to be more similar, relative shear moduli correlated in some but not all regions of the disc. The low number of correlations between relative shear moduli from two loading regimes and between relative shear moduli and relaxation times suggest the limitation of considering different loading conditions, applied to the same disc, as independent. Continuing development of these and similar inverse methods could achieve greater accuracy by constraining multiple sets of deformation data – obtained in orthogonal planes or even through the full tissue volume – to a single, nonhomogeneous model.

In summary, we applied dualMRI to measure the deformation of the disc under cyclic axial compression as well as anterior bending. Deformations and shear modulus, in addition to relaxation times *T_1_* and *T_2_*, were calculated in a coronal and a sagittal plane through the midline of the disc. This work demonstrates the potential for dualMRI in orthogonal planes to collect greater tissue deformation information without the imaging time cost of a full-volume multi-slice acquisition. Our application of inverse modeling to estimate relative shear modulus in the disc enabled investigation of potential correlations to relaxation times *T_1_* and *T_2_*. We found that the estimated shear modulus did not consistently show significant correlations with relaxation times, and that relative shear moduli estimated from different loading regimes may differ. Consequently, we viewed image-based elastography and relaxometry as complementary measures of disc structure and function with potential to assess degeneration in longitudinal studies.

## ACKNOWLEDGEMENTS

The authors gratefully acknowledge the financial support received from NSF grants 1100554 (Neu), 1944394 (Chan), and 2149946 (Chan), and NIH grants R01 AR063712 (Neu), R21 AR064178 (Neu), R25 EB013029-02 (Rundell/Irazoqui, Purdue University), and S10 RR019920-01 (Wyrwicz, Northshore University Healthsystem Research Institute).

