## Supplemental Data for "Intervertebral Disc Elastography to Relate Shear Modulus and Relaxometry in Compression and Bending"

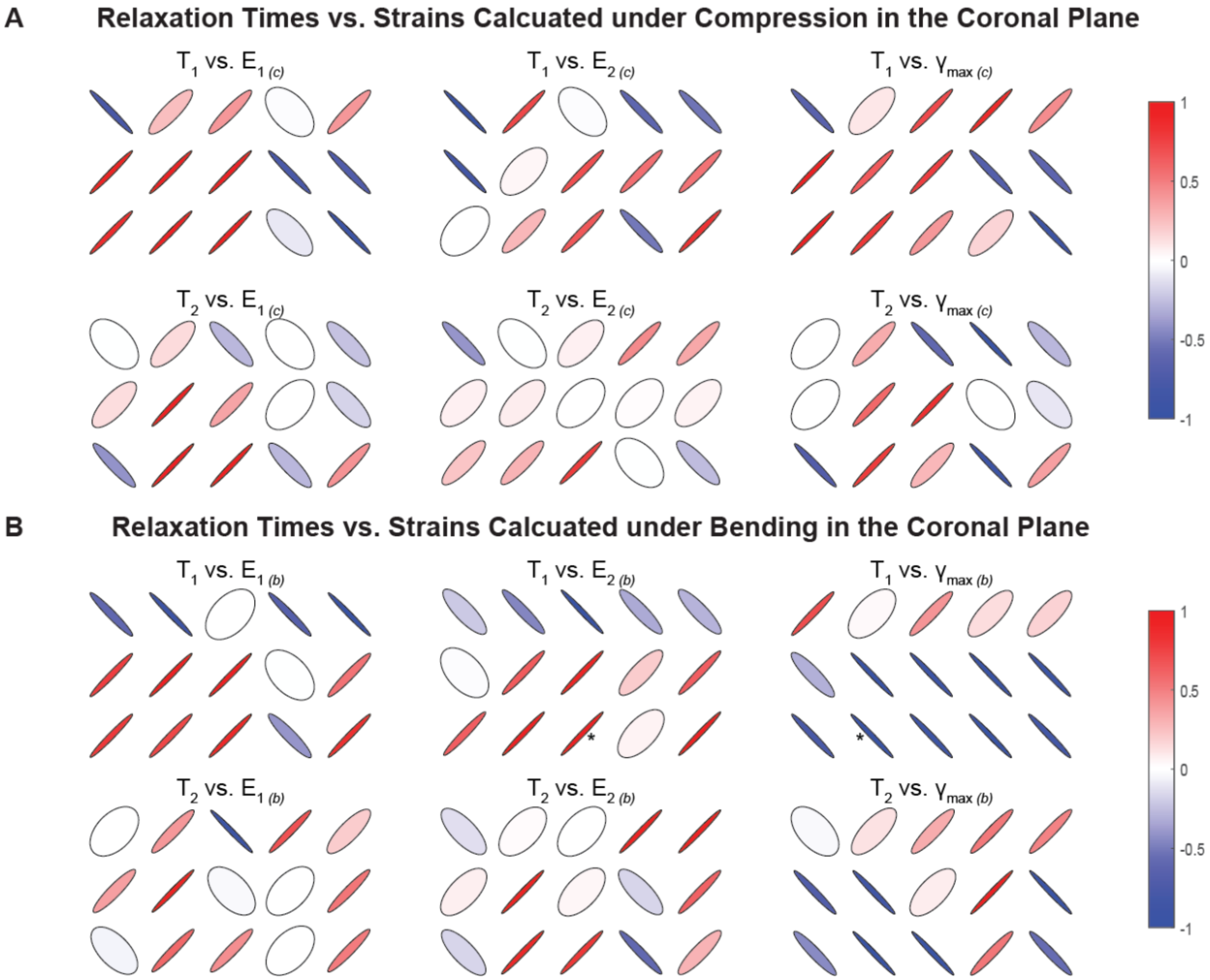


**Supplemental Figure 1.** Correlations by region within the disc between relaxation times and calculated strains from dualMRI in the coronal plane.

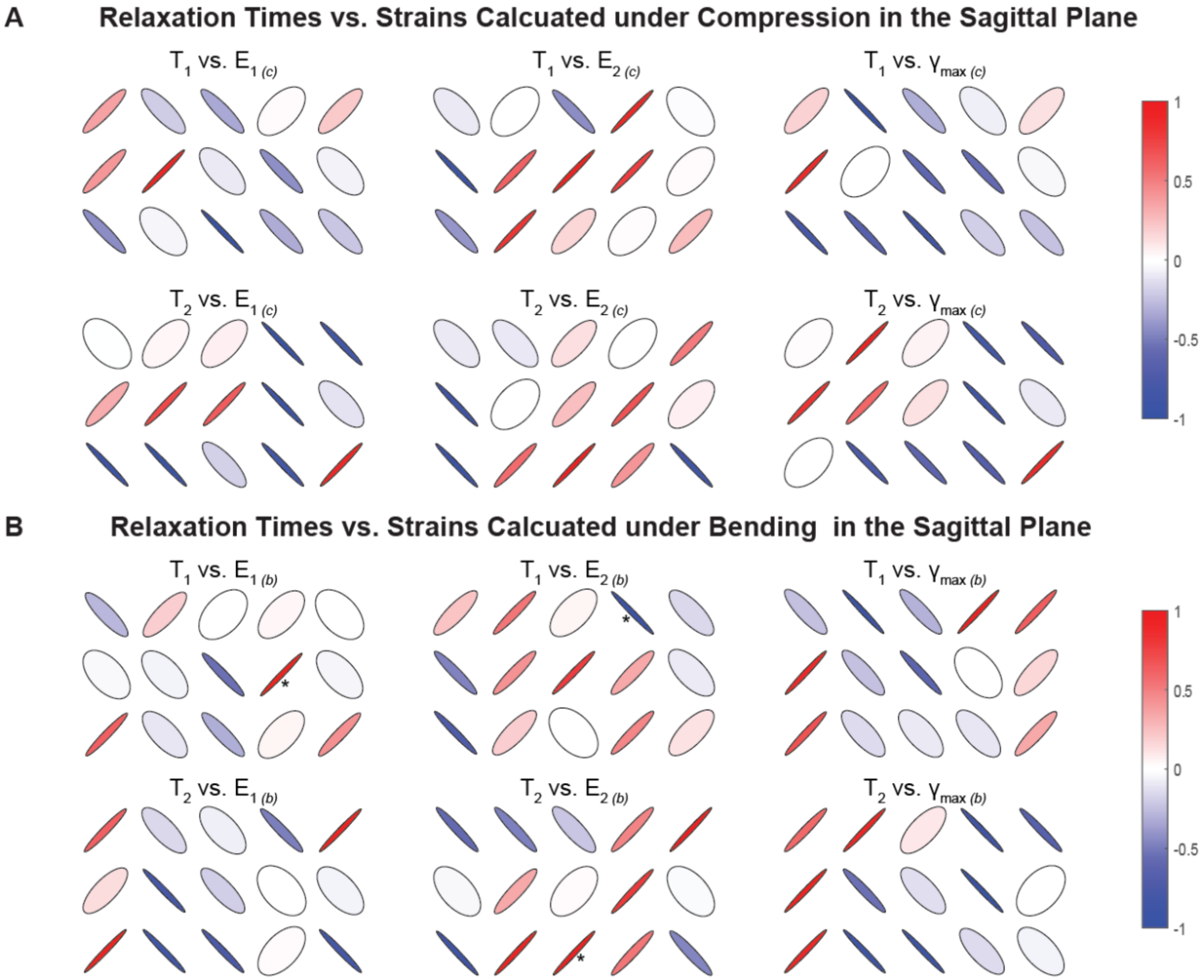


**Supplemental Figure 2.** Correlations by region within the disc between relaxation times and calculated strains from dualMRI in the sagittal plane.
